# ETAP-CLF: an ESM3-based transformer attention framework for binary protein classification

**DOI:** 10.64898/2026.08.16.745129

**Authors:** Jianyu Ren, Hanli Jiang, Puchangxin Li, Xin Yang, Lingwei Mei, Hai Tong, Li Lin

## Abstract

Binary protein classification supports diverse tasks in computational biology, including pathway-membership inference and sequence-based candidate prioritization. Protein language models generate information-rich residue-level representations, but downstream classifiers commonly compress them using fixed pooling operations that may discard task-relevant sequence context. We present ETAP-CLF, a compact framework that combines pretrained per-residue ESM3 embeddings with lightweight transformer contextualization and learned attention pooling to classify variable-length proteins and generate residue-level attention scores. The ESM3 parameters remained frozen, and the same ETAP-CLF architecture and hyperparameter configuration were used across ferroptosis-, senescence-, and pyroptosis-associated protein prediction. ETAP-CLF achieved AUROCs of 0.98, 0.95 and 0.91 for these tasks, respectively. In the ferroptosis benchmark, ETAP-CLF outperformed the evaluated published models. These results demonstrate that a common downstream design can adapt to multiple process-associated classification tasks without fine-tuning the task-specific model architecture. ETAP-CLF provides a generalizable approach for sequence-based protein prioritization and a basis for broader evaluation across binary protein-classification problems.

## Introduction

Sequence-based prediction of protein function and biological-process association is a recurring problem in computational biology, supporting functional annotation, pathway-membership inference and candidate prioritization^1,2^. For individual functions or processes, these questions are often formulated as binary classification tasks; however, proteins vary substantially in length and domain composition, while experimentally supported labels may be limited or incomplete ^3^. Compounding these challenges, sequence homology can inflate performance estimates when closely related proteins occur across training and test partitions. TAPE and PEER helped formalize protein representation learning through standardized downstream tasks and evaluation settings designed to assess biologically meaningful generalization ^4,5^. More generally, rigorous protein machine learning requires evaluation procedures that distinguish recognition of related sequence families from generalization beyond them ^6^.

Self-supervised protein language models have changed how protein sequences are represented for supervised learning. Large-scale pretraining enables these models to encode biochemical, structural and evolutionary information in contextualized residue representations learned from sequence data ^7^. The ESM family has progressively expanded this capability: ESM-2 demonstrated that evolutionary-scale language modelling can support atomic-level structure prediction from individual sequences, whereas ESM3 jointly models protein sequence, structure and function within a multimodal generative framework ^8,9^. When ESM3 is used as a frozen feature extractor, it provides contextualized, per-residue representations without requiring task-specific updates to its pretrained parameters. The effectiveness of this strategy consequently depends on how the downstream model transforms and aggregates those representations.

A protein-level classifier must convert a variable-length sequence of residue embeddings into a fixed-dimensional representation. Mean pooling and special-token pooling are computationally efficient, but these fixed operations cannot learn which residues or sequence regions are most informative for a particular task. Learned aggregation of protein-language-model embeddings is already established: Light Attention assigns trainable weights to frozen residue-level embeddings, and subsequent methods have explored optimal-transport, self-attention and locality-aware pooling ^10–12^. Downstream transformer processing of protein-language-model embeddings has also been used in protein classification and sequence-effect prediction ^13,14^. These studies establish transformer contextualization and attention-based aggregation as useful design patterns rather than unexplored mechanisms. They motivate examining whether a compact ESM3-based implementation can be held constant across multiple biological-process classification tasks while retaining residue-level attention scores for exploratory analysis.

Ferroptosis, cellular senescence and pyroptosis provide related but biologically distinct settings in which to examine such portability. Ferroptosis is an iron-dependent form of regulated cell death driven by phospholipid peroxidation and is implicated in cancer, degenerative disorders and tissue injury ^15^. Cellular senescence is not a cell-death modality but a heterogeneous stress-associated state characterized principally by stable cell-cycle arrest, with context-dependent roles in development, tissue repair, tumour suppression, aging and chronic disease ^16,17^. Pyroptosis is a gasdermin-mediated lytic form of regulated cell death that promotes inflammatory-mediator release and contributes to antimicrobial defence, inflammatory disorders and cancer ^18^. Because association with these processes may depend on cellular context, molecular interactions and regulatory state, sequence-based predictions should be interpreted as candidate-prioritization evidence rather than proof of mechanistic participation.

Among these applications, ferroptosis-associated protein prediction has the most developed sequence-based computational literature. FerrDb and its successive updates provide manually curated collections of ferroptosis regulators, markers and disease associations supported by published evidence^19–21^. Computational prediction has progressed from handcrafted multi-view sequence features in FRP-XGBoost to pooled ESM-2 feature integration in PLM-FRP and contrastive capsule learning in FeroConCap ^22–24^. For cellular senescence, SenSeqNet combined ESM-2 embeddings with a hybrid LSTM-convolutional architecture to predict senescence-associated proteins from sequence ^25^. By contrast, dedicated sequence-based classification of pyroptosis-associated proteins remains comparatively underexplored. Collectively, these developments create an opportunity to evaluate a common residue-level representation and classification workflow across all three processes.

Here, we present ETAP-CLF, a compact framework that combines per-residue ESM3 embeddings with lightweight transformer contextualization, learned attention pooling and binary classification. We use ferroptosis as the primary benchmark, assess performance relative to existing predictors and examine recovery in an external dataset which contains ferroptosis related genes and other regulated cell-death genes. Learned pooling weights are reported as residue-level attention scores for exploratory prioritization, not as evidence of causal or mechanistic importance. ETAP-CLF therefore provides a generalizable framework that can be trained with task-specific labels for diverse binary protein-classification problems.

## Results

### Overview of the ETAP-CLF architecture and evaluation framework

ETAP-CLF was developed as a compact supervised framework that operates directly on per-residue protein-language-model representations (**Figure 1A**). Each protein sequence was encoded into residue-level embeddings by the pretrained ESM3 backbone, whose parameters were held fixed throughout downstream training. ESM3 is a multimodal protein language model pretrained to jointly represent protein sequence, structure and function ^9^. The resulting embeddings were projected into a compact latent space, contextualized using a lightweight four-layer transformer encoder and aggregated by learned single-query attention pooling to generate a protein-level representation for binary classification (**Methods**). Related approaches have previously used trainable attention mechanisms to aggregate frozen residue-level protein-language-model embeddings ^10,12,14^.

This architecture accommodates variable-length proteins while retaining the learned pooling weights as residue-level attention scores. These scores were used for exploratory residue prioritization and were not interpreted as causal or mechanistic attributions. During task-specific training, the ESM3 backbone was held fixed, whereas all downstream ETAP-CLF components, including the embedding projection, transformer encoder, learned attention-pooling module and binary classifier, were updated by backpropagation using positive and negative labels. Together, these downstream components contained approximately 2.50 million trainable parameters. ETAP-CLF was trained separately for ferroptosis-, senescence- and pyroptosis-associated protein prediction, while the architecture and hyperparameter configuration were kept unchanged across tasks (**Figure 1B–C; Methods**). Therefore, the portability analysis evaluated a common modelling framework rather than a single set of classifier weights or a zero-shot model.

The evaluation framework was designed to distinguish performance under conventional random sequence partitioning from generalization to previously unseen genes (**Figure 1B**). Ferroptosis-associated protein prediction served as the primary benchmark. An internal random split represented the evaluation setting used by existing ferroptosis predictors, including FRP-XGBoost, PLM-FRP and FeroConCap ^22–24^. A stricter gene-disjoint external evaluation reserved all sequences from selected ferroptosis-associated and hard-negative regulated-cell-death genes, ensuring that no gene contributed sequences to both model development and external evaluation. Selected published predictors were retrained using the same partitions, allowing the effect of replacing random partitioning with gene-disjoint evaluation to be assessed across different modelling approaches (**Methods**).

The unchanged ETAP-CLF architecture and hyperparameter configuration were subsequently applied to senescence- and pyroptosis-associated protein prediction to evaluate portability across biological-process classification tasks. The senescence analysis used the same sequence benchmark previously developed for SenSeqNet, enabling direct evaluation against a purpose-built senescence-associated protein predictor ^25^. For each task, sequence-level predictions were also aggregated by gene to calculate per-gene recovery rates. Genes represented by sufficient test sequences and consistently recovered by ETAP-CLF were examined using pathway-enrichment analysis (**Figure 1D; Methods**). The complete evaluation framework therefore addressed performance under random partitioning, generalization to genes excluded from training and reuse of the same design across distinct binary protein-classification tasks.

**Figure 1.**
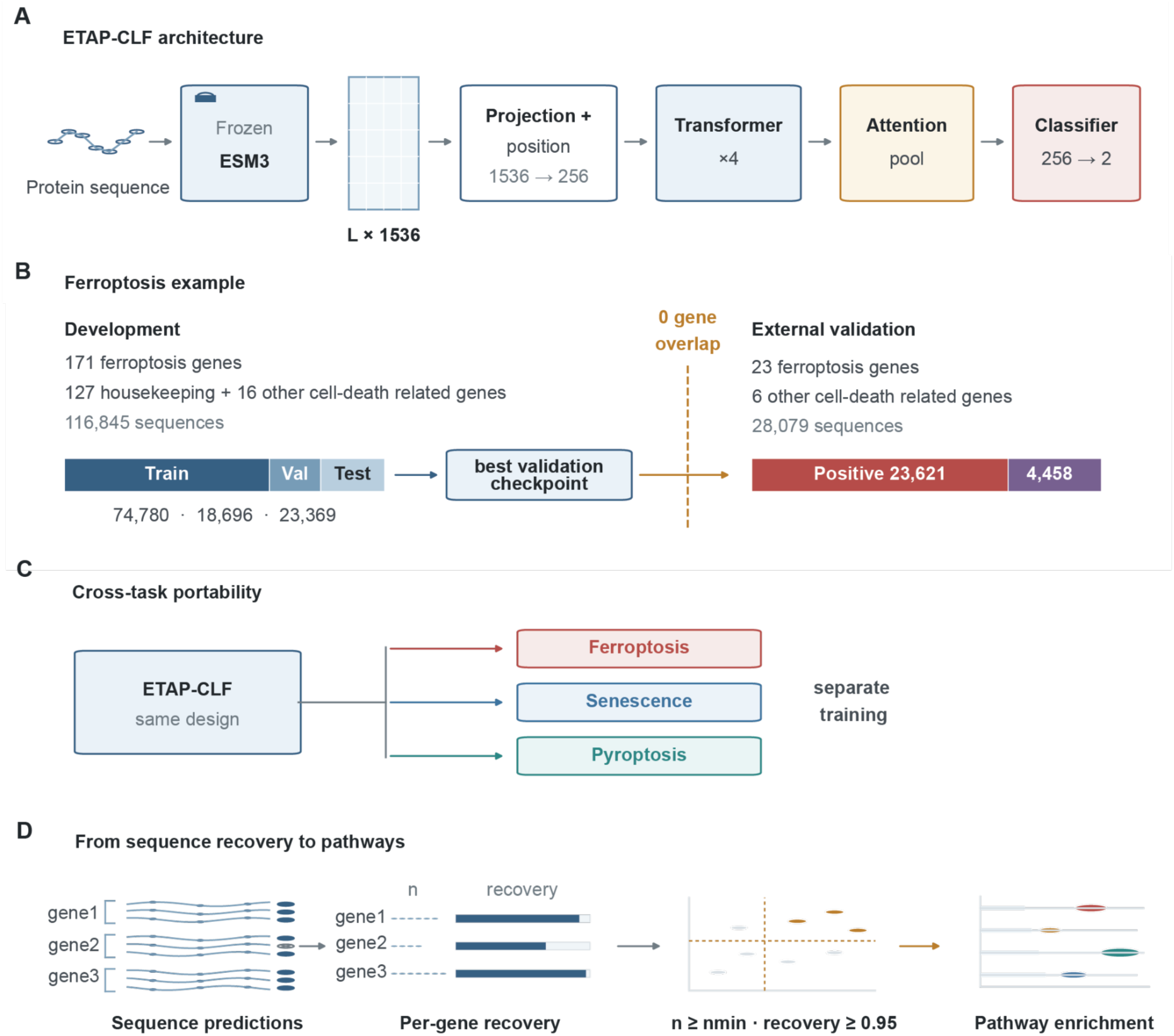
ETAP-CLF architecture and analysis workflow. **(A)** Frozen ESM3 embeddings were projected and processed by four Transformer layers, attention pooling and a binary classifier. **(B)** The ferroptosis development cohort was divided into training, validation and test sets, and the best checkpoint was evaluated on an external cohort with no overlapping genes. **(C)** The same architecture and hyperparameters were used for ferroptosis, senescence and pyroptosis, with separate task-specific training. **(D)** Test predictions were aggregated by gene. Positive genes meeting the sequence-support threshold (n_test ≥ 30 for ferroptosis and senescence; ≥10 for pyroptosis) and recovery ≥0.95 were used for pathway-enrichment analysis.

### Dataset composition and sequence characteristics across the three classification tasks

The final ferroptosis, senescence and pyroptosis datasets differed substantially in scale, class composition and evaluation design (**Table 1**). Ferroptosis provided the primary benchmark, with housekeeping proteins and proteins associated with other regulated cell-death processes serving as negative groups. Senescence represented the largest cluster-representative sequence cohort, whereas the pyroptosis dataset contained both functionally diverse comparison proteins and proteins associated with related regulated cell-death processes. Across all three tasks, sequences were clustered using MMseqs2 at a sequence-identity threshold of 0.30, and one representative sequence per cluster was retained for downstream analysis^26^. This procedure substantially reduced sequence redundancy and the potential for sequence-similarity leakage between data partitions (**Methods**).

The initial ferroptosis collection contained 424,637 ferroptosis-associated and 341,849 housekeeping protein sequences. Following MMseqs2 clustering, 84,335 ferroptosis-associated and 18,276 housekeeping cluster-representative sequences were retained. An additional 18,692 sequences were obtained from 22 genes associated with other regulated cell-death processes. Of these, 14,234 sequences from 16 genes were included in model development, whereas 4,458 sequences from 6 genes were reserved for gene-disjoint external evaluation. The model-development cohort therefore comprised 171 ferroptosis-associated genes, 127 housekeeping genes and 16 genes associated with other regulated cell-death processes. Ferroptosis annotations were obtained from FerrDb V2 ^20^. The gene-disjoint external cohort contained 28,079 sequences from 23 ferroptosis-associated genes and 6 other cell-death-related genes that were entirely excluded from training. These cohorts supported two distinct evaluations: discrimination among sequences derived from represented genes and generalization to sequences from previously unseen genes. Including proteins from related regulated cell-death processes provided a more stringent assessment of specificity than comparison with housekeeping proteins alone.

The initial senescence collection comprised 292,759 senescence-associated and 373,098 comparison protein sequences. MMseqs2 clustering reduced this collection to 43,381 senescence-associated and 32,673 comparison sequences, resulting in a final benchmark of 76,054 cluster representatives. Senescence-associated sequences accounted for approximately 57% of the final dataset. The comparison group represented seven functional protein categories and therefore provided a biologically diverse classification background. The established SenSeqNet benchmark partition was retained, allowing direct comparison with a purpose-built senescence-associated protein predictor ^25^.

The initial pyroptosis collection contained 268,230 pyroptosis-associated sequences, 422,399 sequences representing ten functionally diverse protein categories and 32,095 sequences derived from genes associated with other regulated cell-death processes. Following MMseqs2 clustering, 14,300 pyroptosis-associated sequences, 17,531 functionally diverse comparison and regulated cell-death-related comparison sequences were retained, resulting in a final cohort of 31,831 cluster representatives. Pyroptosis-associated sequences accounted for approximately 45% of this cohort. The dataset represented 177 pyroptosis-associated genes and 187 comparison genes, producing an approximately balanced classification task at both the gene and sequence levels. The comparison groups comprised 165 genes from ten functional protein categories and 22 genes associated with related regulated cell-death processes, including 6 apoptosis-associated, 5 necroptosis-associated and 11 ferroptosis-associated genes. This composition enabled separate evaluation against broadly distributed protein functions and closely related cell-death processes, providing a stringent assessment of pyroptosis-associated protein classification.

**Table 1.** Dataset composition and redundancy control across the three classification tasks.

| Task | Gene counts | Raw Sequences | Representative Sequences | Processing threshold |
| --- | --- | --- | --- | --- |
| Ferroptosis | 194/149 | 424,637/341,849 | 107,956/36,968 | 0.3 |
| Senescence | 210/148 | 292,759/373,098 | 43,381/32,673 | 0.3 |
| Pyroptosis | 177/187 | 268,230/454,494 | 14,300/17,531 | 0.3 |

Note: Gene and sequence counts are reported as positive/negative. For example, 194/149 denotes 194 positive genes and 149 negative genes.

### Transformer contextualization and learned attention pooling produce more class-separable representations

To examine how transformer contextualization and residue aggregation shaped the protein-level representation space, the same ferroptosis internal test sequences were summarized using five strategies: mean-pooled raw ESM3 embeddings, max-pooled raw ESM3 embeddings, mean-pooled transformer-contextualized embeddings, max-pooled transformer-contextualized embeddings and the complete ETAP-CLF representation combining transformer contextualization with learned attention pooling. The resulting representations were visualized using uniform manifold approximation and projection with identical settings across all five strategies ^27^ (**Figure 2A-E; Methods**).

Raw ESM3 representations obtained using mean or max pooling showed substantial overlap between ferroptosis-associated and negative sequences (**Figure 2A,B**). Transformer contextualization improved the organization of both the mean- and max-pooled representations, although the two classes remained partially intermixed (**Figure 2C,D**). The complete ETAP-CLF representation generated using transformer contextualization and learned attention pooling displayed the clearest visual separation between the classes (**Figure 2E**). These findings indicate that transformer contextualization reorganized the frozen ESM3 embeddings into a more label-aligned representation space, while learned attention pooling provided the strongest qualitative class separation among the five evaluated strategies.

**Figure 2.**
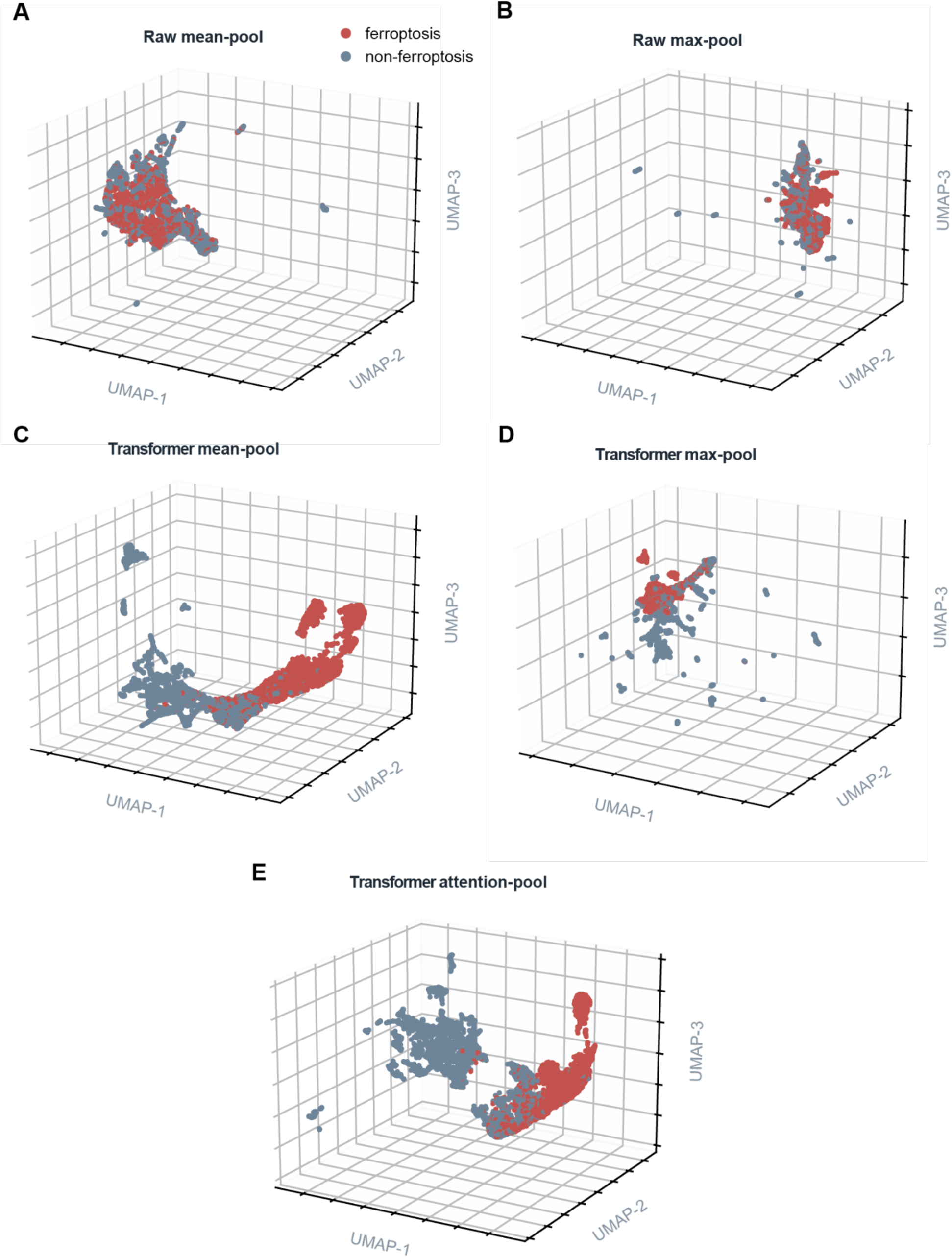
Transformer contextualization and learned attention pooling reshape the ferroptosis protein-representation space. Three-dimensional UMAP projections of protein-level representations from the same internal ferroptosis test sequences. **(A)** Mean-pooled raw ESM3 residue embeddings. **(B)** Max-pooled raw ESM3 residue embeddings. **(C)** Mean-pooled transformer-contextualized embeddings. **(D)** Max-pooled transformer-contextualized embeddings. **(E)** Complete ETAP-CLF representations generated through transformer contextualization and learned attention pooling. Each point represents one protein sequence; red indicates ferroptosis-associated sequences and blue indicates negative sequences. Identical preprocessing and UMAP settings were used across all five representations. Transformer contextualization increased visual class organization, while the complete ETAP-CLF representation displayed the clearest qualitative separation.

### ETAP-CLF achieves strong performance in ferroptosis-associated protein classification

ETAP-CLF showed strong discrimination of ferroptosis-associated proteins in the internal random-split test set of 23,369 sequences (**Figure 3A-B**; **Table 2**). The test set contained 16,870 positive and 6,499 negative sequences. ETAP-CLF achieved an AUROC of 0.980, indicating that ferroptosis-associated sequences generally received higher prediction scores than negative sequences across classification thresholds. Also the model achieved an accuracy of 0.931, a sensitivity of 0.946 and a specificity of 0.893. The model correctly classified 15,960 positive sequences and 5,805 negative sequences, while misclassifying 910 positive sequences and 694 negative sequences. The higher sensitivity than specificity indicated that ETAP-CLF prioritized the recovery of ferroptosis-associated proteins.

These results demonstrate that frozen ESM3 residue representations contain signals that can be adapted effectively by the ETAP-CLF downstream architecture for ferroptosis-associated protein classification. However, because the internal evaluation used a random sequence split, different sequences associated with the same genes could occur across the model-development and test partitions. The strong internal performance therefore reflects discrimination among sequences from represented genes and does not by itself establish generalization to previously unseen genes. This distinction was examined through the subsequent gene-disjoint external evaluation.

### Gene-disjoint external evaluation demonstrates generalization to unseen ferroptosis-associated genes

To assess generalization to previously unseen genes, ETAP-CLF was evaluated on a gene-disjoint external cohort containing 28,079 protein sequences. This cohort comprised 23,621 ferroptosis-associated sequences and 4,458 sequences associated with other regulated cell-death processes. All genes represented in the external cohort were excluded from model development. Housekeeping proteins were not included in this evaluation, allowing specificity to be assessed against proteins involved in biologically related cell-death pathways (**Figure 3C**).

ETAP-CLF achieved an AUROC of 0.801, an accuracy of 0.765, a sensitivity of 0.791 and a specificity of 0.629 (**Table 2**). Thus, the model recovered 79.1% of ferroptosis-associated sequences while correctly rejecting 62.9% of sequences associated with other regulated cell-death processes. These results demonstrate that ETAP-CLF retained meaningful discriminatory ability when applied to sequences from genes not encountered during training. Nevertheless, the lower specificity relative to sensitivity indicates that distinguishing ferroptosis-associated proteins from proteins participating in closely related cell-death pathways remains challenging.

### ETAP-CLF outperforms published ferroptosis predictors under matched evaluation conditions

To minimize differences arising from datasets and evaluation procedures, FRP-XGBoost, PLM-FRP and FeroConCap were retrained using the same model-development sequences as ETAP-CLF and evaluated on identical internal and gene-disjoint external test cohorts (**Methods**). FRP-XGBoost uses handcrafted sequence descriptors with XGBoost, whereas PLM-FRP combines mean-pooled ESM-2 embeddings with sequence-derived features and XGBoost ^22,23^. FeroConCap transforms protein sequences into fractal chaos game representation images and applies capsule networks with supervised contrastive learning ^24^.

On the internal test set (*n* = 23,369), ETAP-CLF achieved the highest AUROC of 0.980, compared with 0.956 for PLM-FRP, 0.912 for FRP-XGBoost and 0.794 for FeroConCap (**Figure 3D**; **Table 2**). These values corresponded to absolute improvements of 0.024, 0.068 and 0.186, respectively. On the gene-disjoint external cohort (*n* = 28,079), ETAP-CLF retained the highest AUROC at 0.801, compared with 0.784 for PLM-FRP, 0.621 for FRP-XGBoost and 0.482 for FeroConCap. Although AUROC decreased for every model under gene-disjoint evaluation, ETAP-CLF showed the highest observed ranking performance in both settings. Because all models were trained and evaluated using the same sequences, labels and partitions, these differences were not attributable to variation in dataset composition.

**Figure 3.**
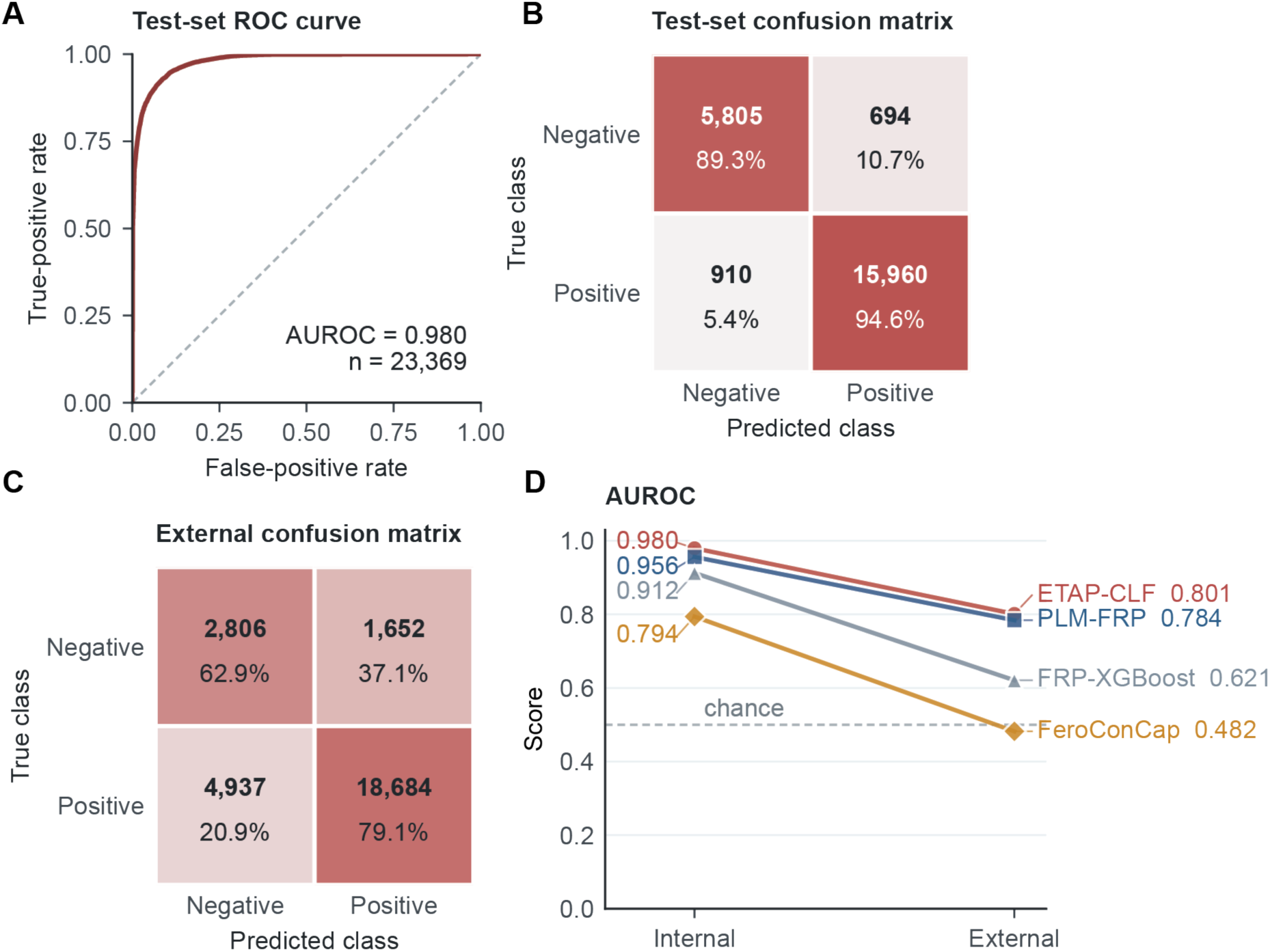
Ferroptosis classification performance and matched comparison with published predictors. **(A)** ROC curve for ETAP-CLF on the internal test set. **(B)** Sequence-level confusion matrix for the internal test set. **(C)** Sequence-level confusion matrix for the gene-disjoint external validation cohort. **(D)** AUROC comparison of ETAP-CLF with published ferroptosis predictors on the matched internal and external benchmark, for which all models were trained and evaluated using the same sequence dataset and data partitions.

**Table 2.** Performance of ETAP-CLF and evaluated benchmark models for ferroptosis.

| Test/External | Model | AUROC | Accuracy | Sensitivity | Specificity |
| --- | --- | --- | --- | --- | --- |
| Test | **ETAP-CLF** | **0.980** | **0.931** | **0.946** | **0.893** |
| Test | PLM-FRP (2025) | 0.956 | 0.909 | 0.957 | 0.784 |
| Test | FRP-XGBoost (2024) | 0.912 | 0.874 | 0.961 | 0.623 |
| Test | FeroConCap (2026) | 0.794 | 0.774 | 0.882 | 0.554 |
| External | **ETAP-CLF** | **0.801** | **0.765** | **0.791** | **0.629** |
| External | PLM-FRP (2025) | 0.784 | 0.716 | 0.713 | 0.734 |
| External | FRP-XGBoost (2024) | 0.621 | 0.834 | 0.932 | 0.316 |
| External | FeroConCap (2026) | 0.482 | 0.528 | 0.560 | 0.354 |

### An unchanged ETAP-CLF configuration supports senescence- and pyroptosis-associated protein prediction

To evaluate the portability of ETAP-CLF, the architecture and hyperparameter configuration used for ferroptosis classification were retained for the senescence and pyroptosis tasks. The pretrained ESM3 backbone remained frozen, whereas separate ETAP-CLF downstream parameters were trained using the labels for each task. Thus, no task-specific architectural or hyperparameter modifications were introduced (**Methods**).

On the held-out senescence test set of 15,211 sequences, ETAP-CLF achieved an AUROC of 0.949, an accuracy of 0.874, a sensitivity of 0.911, and a specificity of 0.823 (**Figure 4A–B**; **Table 3**). These results modestly exceeded the reported performance of the purpose-built SenSeqNet model, which achieved an AUROC of 0.940, an accuracy of 0.864, a sensitivity of 0.905, and a specificity of 0.810 ^25^. Thus, even without fine-tuning the ESM3 backbone or introducing task-specific modifications to the downstream architecture and hyperparameter configuration, ETAP-CLF outperformed the reported SenSeqNet benchmark across all four metrics.

To our knowledge, no dedicated method has previously been reported for classifying pyroptosis-associated proteins directly from primary amino-acid sequences. ETAP-CLF achieved an AUROC of 0.91 in the pyroptosis sequence-classification task, demonstrating that the unchanged configuration captured predictive sequence information associated with pyroptosis (**Figure 4C–D**; **Table 3**). Although this performance was lower than that observed for ferroptosis and senescence, the result supports the feasibility of predicting pyroptosis-associated proteins from primary sequence alone. The lower AUROC may reflect greater sequence heterogeneity among pyroptosis-associated proteins, biological overlap with other regulated cell-death pathways or a stronger dependence of pyroptosis association on cellular context and protein interactions. However, because the three datasets differed in composition and evaluation design, their AUROCs should not be interpreted as direct measurements of their relative biological difficulty.

Collectively, these results show that the same ETAP-CLF architecture and hyperparameter configuration can be trained across multiple process-associated protein-classification tasks without fine-tuning the model. The framework approached the performance of a purpose-built senescence predictor and provided a sequence-based benchmark for pyroptosis-associated protein prediction, supporting its use as a reusable downstream design rather than a task-specific classifier.

**Table 3.** Performance of ETAP-CLF for senescence and pyroptosis classification, with SenSeqNet as the senescence benchmark.

| Task | Model | AUROC | Accuracy | Sensitivity | Specificity |
| --- | --- | --- | --- | --- | --- |
| Senescence | **ETAP-CLF** | **0.949** | **0.874** | **0.911** | **0.824** |
| Senescence | SenSeqNet | 0.940 | 0.864 | 0.905 | 0.810 |
| Pyroptosis | **ETAP-CLF** | **0.909** | **0.821** | **0.785** | **0.850** |

**Figure 4.**
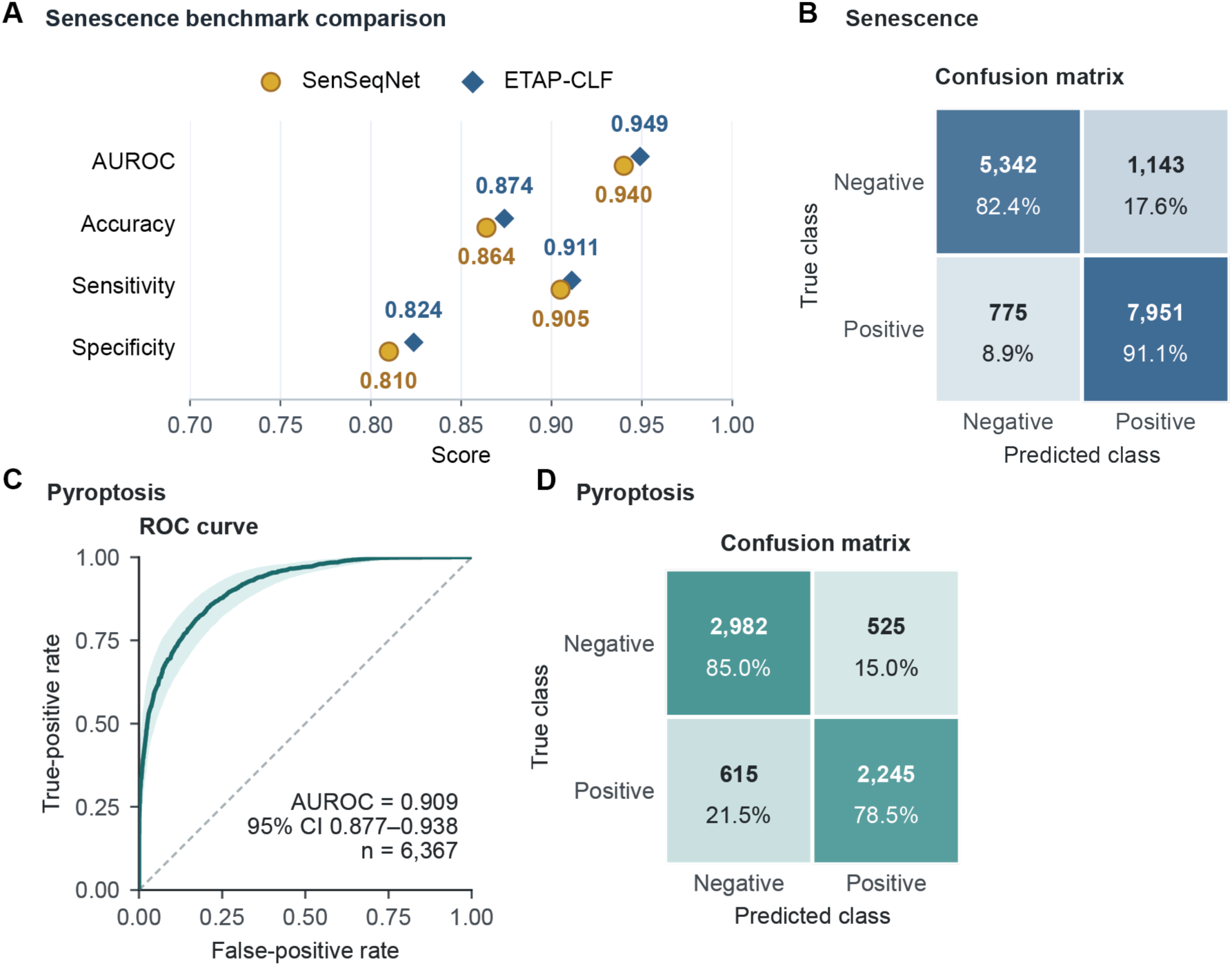
Portability of the unchanged ETAP-CLF configuration to senescence and pyroptosis classification. (**A**) Contextual comparison between ETAP-CLF and the purpose-built SenSeqNet model. (**B**) Confusion matrix for ETAP-CLF to senescence classification. (**C**) AUROC curve and (**D**) confusion matrix for the pyroptosis test set. ESM3 remained frozen, and the same ETAP-CLF architecture and hyperparameter configuration were used for both tasks.

### Consistently recovered genes recapitulate process-relevant biological pathways

To determine whether ETAP-CLF consistently recovered biologically coherent subsets of positive genes, positive-class test predictions were aggregated by gene and summarized as per-gene recovery rates. For ferroptosis and senescence, genes were retained when they were represented by at least 30 test sequences and achieved a recovery rate of at least 0.95. Applying the same sequence threshold to pyroptosis retained only one gene; therefore, the pyroptosis analysis required at least 10 test sequences while retaining the 0.95 recovery-rate threshold. These criteria selected 25 of 169 ferroptosis-associated genes, 17 of 201 senescence-associated genes and 9 of 141 pyroptosis-associated genes represented in the corresponding per-gene test summaries (**Figure 5A–C**). The selected genes represented 13,262, 1,889 and 233 test sequences, respectively, with sequence-weighted recovery rates of 0.980, 0.972 and 0.983. The minimum Wilson 95% lower confidence bounds were 0.838, 0.833 and 0.722, respectively, reflecting the smaller per-gene test counts in the pyroptosis dataset.

The ferroptosis gene set showed the clearest agreement with direct pathway annotations across databases (**Figure 5D**). Eight selected genes, including *ATG7*, *CYBB*, *HSPB1*, *NOX1*, *POR*, *SAT1*, *TF* and *TP53*, overlapped the WikiPathways ferroptosis pathway (48.53-fold enrichment; FDR = 1.41 × 10⁻¹⁰). Four genes also overlapped the KEGG ferroptosis pathway (43.96-fold enrichment; FDR = 3.84 × 10⁻⁵). Mechanistically related enrichment included the KEGG peroxisome pathway (27.14-fold; FDR = 3.61 × 10⁻⁵), cellular response to oxidative stress (24.55-fold; FDR = 3.25 × 10⁻⁵) and p53 signalling (24.03-fold; FDR = 2.30 × 10⁻⁴). Ether-lipid biosynthesis, macroautophagy and NADPH-oxidase activation were also recovered, indicating that the selected genes represented multiple annotated components of ferroptosis-associated lipid, redox and stress-response biology.

The senescence result was distributed across direct senescence annotations and related growth-arrest, stress and inflammatory processes (**Figure 5E**). The selected set was enriched for KEGG cellular senescence through *CXCL8* and *HRAS* (9.45-fold; FDR = 0.0258). The corresponding Reactome cellular-senescence term contained *CXCL8* and *HIRA* but was borderline after correction (9.98-fold; FDR = 0.0508). Stronger linked signals included negative regulation of cell-population proliferation (FDR = 0.0026), Notch signalling (FDR = 0.0030), inflammatory response (FDR = 0.0072), cellular responses to stress (FDR = 0.0074) and the p53 transcriptional network (FDR = 0.0207). Thus, the selected genes recapitulated several characteristic components of senescence biology, although each direct senescence term was supported by only two genes.

The pyroptosis gene set recovered a coherent inflammasome–interleukin-1β axis (**Figure 5F**). Six selected genes overlapped regulation of interleukin-1β production (122.02-fold; FDR = 1.20 × 10⁻¹⁰), and *CASP8*, *NAIP* and *NLRP3* overlapped the direct GO pyroptotic inflammatory-response term (167.28-fold; FDR = 9.70 × 10⁻⁶). Additional enrichment included the Reactome CLEC7A inflammasome pathway (FDR = 3.30 × 10⁻⁵), regulation of NLRP3 inflammasome assembly (FDR = 3.57 × 10⁻⁵), KEGG NOD-like receptor signalling (FDR = 2.39 × 10⁻⁴) and positive regulation of inflammasome-mediated signalling (FDR = 5.24 × 10⁻⁴). The exact Reactome Pyroptosis and Inflammasomes terms each contained fewer than two selected genes and were therefore not tested.

Collectively, the consistently recovered genes mapped to direct or mechanistically related annotations for each target process. Because the input genes were already curated as process associated and enrichment was evaluated against pathway-wide annotated gene universes, these results constitute a biological-coherence analysis rather than independent discovery of pathway membership.

**Figure 5.**
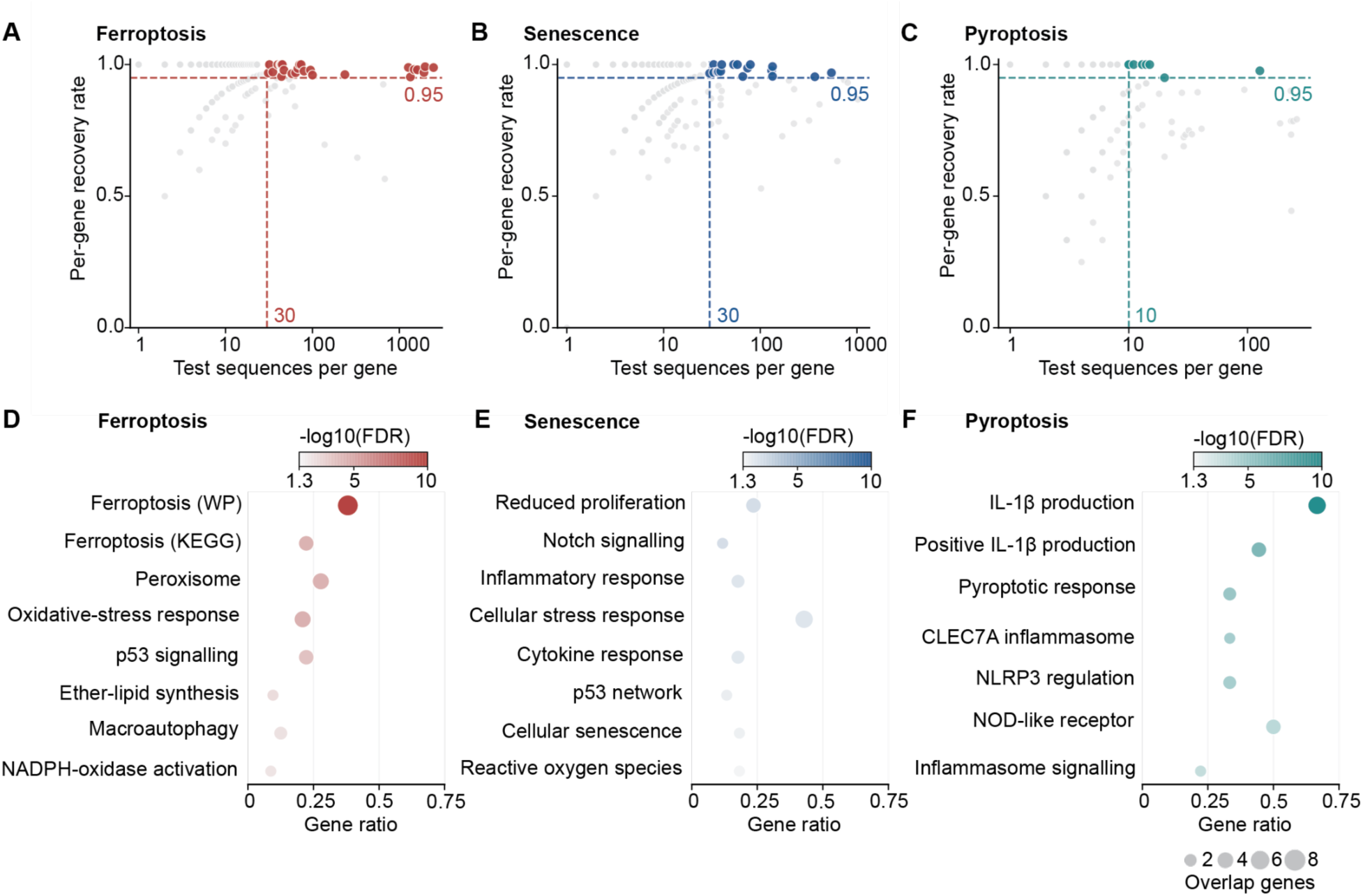
Consistently recovered genes represent process-relevant biological pathways. a–c, Per-gene recovery rate versus the number of held-out positive sequences for ferroptosis (**A**), senescence (**B**) and pyroptosis (**C**). Coloured points met the selection criteria: n ≥ 30 and recovery ≥ 0.95 for ferroptosis and senescence, and n ≥ 10 and recovery ≥ 0.95 for pyroptosis. These criteria retained 25 of 169, 17 of 201 and 9 of 141 genes, with pooled recovery rates of 0.980, 0.972 and 0.983, respectively. (**D–F)** Pathway enrichment among the selected genes. Gene ratio is shown on the x axis, point size represents the number of overlapping genes and colour indicates −log10(FDR). One-sided hypergeometric tests used each pathway library’s annotated gene union as the background, followed by library-specific Benjamini– Hochberg correction. Panels show selected terms with FDR < 0.05; complete results are provided in the source data. Recovery rates are descriptive sequence-level estimates.

## Discussion

This study evaluated whether frozen residue-level ESM3 representations could support multiple biological-process classification tasks through a compact downstream architecture. ETAP-CLF combined transformer contextualisation with learned attention pooling and achieved strong performance for ferroptosis-, senescence- and pyroptosis-associated protein prediction. The same architecture and hyperparameters were retained across tasks, although separate downstream weights were trained for each dataset. ETAP-CLF should therefore be considered a reusable supervised framework rather than a zero-shot classifier.

Protein language models encode evolutionary, structural and functional information within residue-level representations ^7,9^, but their use for protein-level prediction depends on how these variable-length representations are aggregated. Fixed mean or max pooling may discard task-relevant sequence information, whereas learned pooling can prioritise informative residues or regions ^10,12^. Consistent with this interpretation, transformer contextualisation improved the qualitative organisation of the ferroptosis representation space, and the complete attention-pooled representation showed the clearest UMAP class separation. However, UMAP remains descriptive and does not independently establish the contribution of each model component. Similarly, attention weights should be interpreted as exploratory residue-prioritisation scores rather than causal explanations.

The matched ferroptosis comparison reduced the confounding effects of using different datasets and evaluation procedures. On the internal test set, ETAP-CLF achieved an AUROC of 0.980, compared with 0.956 for PLM-FRP, 0.912 for FRP-XGBoost and 0.794 for FeroConCap. On the gene-disjoint cohort, the corresponding AUROCs were 0.801, 0.784, 0.621 and 0.482. ETAP-CLF therefore had the highest observed ranking performance under both evaluation settings, although its external advantage over PLM-FRP was modest and should not be interpreted as statistically significant without paired uncertainty estimates. The decline observed for every model under gene-disjoint evaluation also illustrates how random sequence partitions can produce optimistic estimates when sequences from represented genes occur across model-development and test sets ^6,28^. Nevertheless, the external cohort also replaced housekeeping proteins with proteins involved in related cell-death processes; therefore, the performance reduction cannot be attributed to gene disjointness alone. ETAP-CLF retained meaningful discrimination, but its sensitivity of 0.791 and specificity of 0.629 show that distinguishing ferroptosis-associated proteins from biologically related cell-death proteins remains challenging.

The portability analyses further supported the use of ETAP-CLF as a common downstream design. For senescence, ETAP-CLF achieved an AUROC of 0.949, within the same performance range as the purpose-built SenSeqNet model ^25^. For pyroptosis, ETAP-CLF achieved an AUROC of 0.909, providing an initial benchmark for direct sequence-based classification in this comparatively underexplored setting. Performance differences among the three tasks should not be interpreted as differences in intrinsic biological difficulty because their datasets and comparison groups were not directly equivalent. Moreover, ferroptosis, senescence and pyroptosis depend on cellular context, molecular interactions and regulatory state in addition to primary sequence ^15,16,18^. Predictions should therefore be viewed as evidence for candidate prioritisation rather than proof of mechanistic involvement.

Consistently recovered genes were enriched for pathways coherent with each target process, including lipid-peroxidation and oxidative-stress pathways for ferroptosis, growth-arrest and inflammatory programmes for senescence, and inflammasome–interleukin-1β signalling for pyroptosis. These results support the biological coherence of the model predictions but do not constitute independent pathway validation because the analysed genes were already annotated as process associated and were selected according to model recovery.

Several limitations remain. Gene-level labels were propagated to multiple protein sequences even though isoforms, orthologues and fragments may differ biologically, and incomplete annotations can introduce uncertainty into both positive and comparison classes. Thirty-percent sequence clustering reduced redundancy but did not guarantee family- or structure-disjoint evaluation. In addition, the study used one frozen ESM3 checkpoint, excluded proteins exceeding its supported length and did not experimentally validate the residue-level attention scores or prioritised proteins. Future work should include repeated training runs, paired confidence intervals, calibrated prediction thresholds and gene-, family-, species- and structure-disjoint benchmarks. Within these limitations, ETAP-CLF demonstrates that compact contextualisation and learned aggregation can adapt frozen ESM3 representations to multiple binary protein-classification tasks while highlighting the importance of realistic generalisation assessment.

## Methods

### Study design

ETAP-CLF was evaluated as a reusable downstream architecture for binary protein classification. For every task, protein sequences were encoded using the same pretrained ESM3 checkpoint, and a separate ETAP-CLF model was trained using task-specific labels. ESM3 remained frozen, and the ETAP-CLF architecture and downstream hyperparameter configuration were held constant across ferroptosis-, senescence- and pyroptosis-associated protein prediction; only the learned downstream weights differed among tasks.

Ferroptosis served as the primary benchmark. The evaluation included an internal stratified sequence-level partition and a gene-disjoint cohort in which every sequence from the selected external genes was excluded from model development. Senescence and pyroptosis were used to test architectural portability without task-specific modification.

### Gene sets and protein-sequence acquisition

Ferroptosis-associated genes were obtained from FerrDb V2 ^20^. The development roster contained 163 drivers and 8 markers. Functional comparison genes represented ten broad protein categories, and 22 genes associated with apoptosis, necroptosis or pyroptosis were included as hard negatives (**Supplementary Tables S1, S2 and S5**).

The senescence benchmark followed SenSeqNet ^25^ and comprised 163 CellAge entries and 47 genes supported by at least two in vivo senescence resources, together with 148 comparison entries from seven functional categories ^29^. The pyroptosis collection was derived from one Reactome and six Gene Ontology Biological Process gene sets. Sequence records were available for 177 pyroptosis-associated genes. The comparison set included ten functional groups and 22 hard-negative genes associated with apoptosis, necroptosis or ferroptosis (**Supplementary Tables S3–S5**).

Ferroptosis and pyroptosis sequences were obtained from UniParc, whereas the senescence sequences were inherited from the UniProtKB-based SenSeqNet benchmark ^25,30^.

### Sequence quality control and redundancy reduction

Sequences shorter than 10 amino acids were removed. Redundancy was reduced with MMseqs2 by clustering at a minimum sequence-identity threshold of 0.30 and retaining one representative sequence per cluster.

The representative sequences were then filtered for ESM3 compatibility before embedding generation: entries containing residues unsupported by the ESM3 tokenizer, and proteins exceeding 2,048 residues, were excluded. In the ferroptosis-associated and housekeeping collection, this step removed one unsupported-residue sequence and 1,349 overlength sequences, leaving 102,611 sequences. Together with 14,234 hard-negative sequences from 16 regulated-cell-death genes, 116,845 ESM3-compatible sequences entered ferroptosis model development **(Supplementary Table S5)**.

Cluster-representative selection reduces redundancy but does not eliminate gene-level dependence, because different sequences assigned to the same gene can be distributed across internal partitions. The gene-disjoint cohort was therefore used to assess generalization to genes excluded from model development.

### Ferroptosis model-development and gene-disjoint cohorts

After ESM3 compatibility filtering, the ferroptosis-associated and housekeeping collection contained 102,611 sequences. A further 14,234 sequences from 16 regulated-cell-death genes were included in model development. The resulting 116,845 sequences were divided by stratified random sampling into 74,780 training, 18,696 validation and 23,369 internal test sequences.

The gene-disjoint cohort contained 28,079 sequences: 23,621 sequences from 23 ferroptosis-associated genes and 4,458 sequences from six genes associated with other regulated cell-death processes **(Supplementary Table S2)**. All sequences from these 29 genes were excluded from training, validation, model selection and hyperparameter development. Housekeeping sequences were not included in this evaluation.

### Senescence cohort and partitioning

The SenSeqNet dataset initially contained 292,759 senescence-associated sequences and 373,098 comparison sequences. Clustering with MMseqs2 at 30% sequence identity retained 43,381 positive and 32,673 negative cluster representatives. These sequences were partitioned into training and test sets at an 80:20 ratio using the same random seed reported for SenSeqNet ^25^.

### Pyroptosis cohort and partitioning

The initial pyroptosis collection contained 268,230 pyroptosis-associated sequences, 422,399 sequences from ten functional comparison categories and 32,095 sequences associated with related regulated cell-death processes. After clustering with MMseqs2 at 30% sequence identity, the retained training set comprised 14,300 positive and 17,531 negative sequences, which were divided by stratified random sampling into 20,371 training, 5,093 validation and 6,367 internal test sequences.

### Frozen ESM3 residue embeddings

Residue-level embeddings were generated using ‘esm3_sm_open_v1’, the open small ESM3 checkpoint. ESM3 jointly models protein sequence, structure and function^9^. The checkpoint was placed in evaluation mode, excluded from gradient computation and never updated during any downstream task.

For a protein of length Lᵢ, removal of special tokens produced an embedding matrix 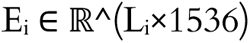. Inference used bfloat16 precision, and cached residue embeddings were stored in HDF5 using float16 values.

### ETAP-CLF architecture

Each 1,536-dimensional residue embedding was projected into a 256-dimensional latent space by a learned linear transformation followed by layer normalization. Sinusoidal positional encodings were added to retain residue order. The projected sequence was processed by four pre-layer-normalized transformer encoder layers, each containing eight attention heads, a 512-dimensional feed-forward module, GELU activation and dropout of 0.1. Padding masks prevented padded positions from contributing to self-attention. This encoder follows the transformer design introduced by Vaswani and colleagues ^31^.

A learned query vector 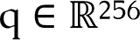 assigned a score to each valid transformer output h_il_:

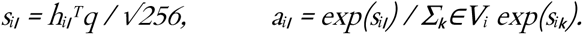

Here, V_i_ denotes the valid, non-padded residue positions. The fixed-dimensional protein representation was the attention-weighted sum:

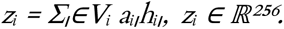

The pooled representation was passed through layer normalization, dropout of 0.1 and a linear layer producing two class logits. The projection, transformer, attention-pooling and classification modules contained 2,503,682 trainable parameters. This count excludes the frozen ESM3 backbone.

### Training and model selection

Separate ETAP-CLF models were trained for the three tasks using identical hyperparameters. Training used unweighted cross-entropy loss and AdamW with a learning rate of 3 × 10⁻⁴, weight decay of 1 × 10⁻⁴ and batch size 64 ^32^. Variable-length sequences were padded within batches, length-bucketed sampling reduced padding overhead and mixed-precision training was used on CUDA-enabled hardware.

The learning rate was increased linearly from 0.01 of its target value during the first three epochs and then reduced by cosine annealing to 1 × 10⁻⁶. Training continued for at most 30 epochs, with early stopping after ten epochs without improvement in validation AUROC. The checkpoint attaining the highest validation AUROC was retained, and test predictions were generated only after model selection.

### Residue-level attention scores

Attention-pooling weights were extracted for valid residues and normalized to sum to one within each protein. They indicate the relative emphasis assigned by the learned pooling layer when constructing the protein representation. These values were treated as exploratory residue-prioritization scores, not causal explanations of residue function, because attention weights are not necessarily faithful explanations of model decisions ^33^.

### Pooling comparison, representation visualization and ablation

To examine the effects of transformer contextualization and residue aggregation, protein-level representations were extracted from the same ferroptosis internal test sequences using five strategies: (i) mean pooling of raw frozen ESM3 residue embeddings, (ii) max pooling of the same embeddings, (iii) mean pooling of transformer-contextualized embeddings, (iv) max pooling of transformer-contextualized embeddings and (v) the complete ETAP-CLF representation obtained by applying learned attention pooling to the transformer-contextualized embeddings. Pooling was restricted to valid residue positions, excluding special tokens and padding.

Each representation matrix was projected independently into three dimensions using UMAP with identical hyperparameters and random seed across all five strategies^27^. The projections were used for qualitative comparison of class organization only. Comparisons between raw and transformer-contextualized representations illustrated the effect of contextualization under the same fixed pooling operation, whereas comparison with the complete ETAP-CLF representation examined the additional effect of learned aggregation. Because UMAP is a nonlinear visualization method, these projections were not used as quantitative evidence that an individual component caused improved classification performance.

### Matched comparison with published ferroptosis predictors

For a dataset-controlled comparison, FRP-XGBoost, PLM-FRP, and FeroConCap were reconstructed and trained on the same development partitions used for ETAP-CLF, with model selection restricted to the validation partition and final evaluation performed on common sequence identifiers. FRP-XGBoost combines amino-acid composition, composition of k-spaced amino-acid pairs, dipeptide deviation from expected mean and grouped tripeptide composition with feature selection and XGBoos ^22^.

PLM-FRP combines 400 DDE features with 1,280-dimensional mean-pooled ESM-2 embeddings and an XGBoost classifier ^8,23^. The present implementation ranked features by XGBoost gain and retained 494 features. FRP-XGBoost retained 299 features after training-only feature ranking and incremental selection.

FeroConCap encodes each protein sequence as a frequency chaos game representation and classifies it with a capsule network employing dynamic routing, optimized under a margin loss with auxiliary reconstruction and supervised-contrastive objectives ^24^. The authors’ published implementation was reused without architectural modification, with gradient clipping added to stabilize training on the present dataset.

### Performance evaluation

The process-associated class was treated as positive for each task. Sequence-level performance was summarized using AUROC, average precision, accuracy, sensitivity, specificity, precision, F1 score and Matthews correlation coefficient. Binary metrics used a probability threshold of 0.5. Confusion matrices and class prevalence were reported with threshold-dependent metrics.

### Gene selection and pathway-enrichment analysis

Positive-class test predictions were aggregated by gene. For each gene g, the per-gene recovery rate was calculated as:

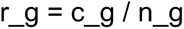

where n_g is the number of positive test sequences assigned to gene g and c_g is the number classified as positive at a probability threshold of 0.5. The per-gene recovery rate therefore corresponds to sensitivity among positive sequences and should not be interpreted as accuracy across both classes. When only n_g and r_g were available, the number of recovered sequences was reconstructed as:

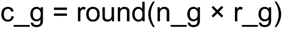

Wilson 95% binomial confidence intervals were calculated to quantify the uncertainty arising from unequal numbers of sequences across genes.

For the ferroptosis and senescence analyses, consistently recovered genes were defined as those satisfying n_g ≥ 30 and r_g ≥ 0.95. Because only one pyroptosis-associated gene met this sequence-count threshold, the pyroptosis analysis used n_g ≥ 10 and r_g ≥ 0.95. The effects of varying the minimum sequence count were evaluated using nearby thresholds and reported in the source data. These filters retained 25 ferroptosis-associated genes, 17 senescence-associated genes and nine pyroptosis-associated genes for pathway analysis.

Pathway over-representation analysis was conducted locally using GMT snapshots labelled KEGG 2026, Reactome Pathways 2024, WikiPathways Human 2024 and Gene Ontology Biological Process 2026 ^34–37^. Gene symbols were standardized to uppercase. For each library, the union of all annotated genes was used as the statistical universe. Query genes absent from this universe were excluded from the analysis for that library. Terms overlapping fewer than two query genes were not tested.

For each pathway-enrichment test:

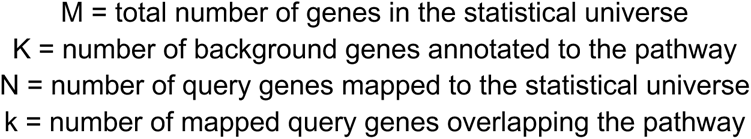

The upper-tail hypergeometric probability of observing at least k overlapping genes was calculated as:

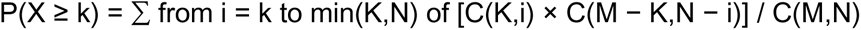

where C(a,b) denotes the number of combinations of b elements selected from a elements. Fold enrichment was calculated as:

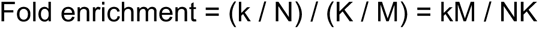

P values were adjusted separately within each pathway library using the Benjamini–Hochberg procedure, and an FDR below 0.05 was considered statistically significant.

## Data availability

The processed sequence datasets and source data underlying the reported results will be deposited in a public repository upon acceptance of this manuscript for publication. During peer review, these materials will be made available confidentially to editors and reviewers upon request. The original data sources are publicly accessible as described in the Methods.

## Code availability

All code used for data preprocessing, model training, evaluation, downstream analysis and visualization, together with documentation for reproducing the principal analyses and figures, will be released through a public GitHub repository upon acceptance of this manuscript for publication. During peer review, access to the private repository will be provided to editors and reviewers upon request.

## Author Contributions

J.R. and H.J. developed the computational model, analysed the data, and prepared the figures. H.J., J.R. and L.L. wrote the manuscript. P.L., X.Y., L.M. and H.T. contributed to data curation, analyses and interpretation. H.J. and L.L conceptualized the idea, supervised the project. L.L. acquired funding. All coauthors reviewed, edited and approved the final manuscript.

## References

1. Bonetta, R. & Valentino, G. Machine learning techniques for protein function prediction. Proteins 88, 397–413 (2020).

2. Radivojac, P. et al. A large-scale evaluation of computational protein function prediction. Nat. Methods 10, 221–227 (2013).

3. Kahanda, I., Funk, C. S., Ullah, F., Verspoor, K. M. & Ben-Hur, A. A close look at protein function prediction evaluation protocols. Gigascience 4, 41 (2015).

4. Xu, M. et al. PEER: A comprehensive and multi-task benchmark for protein sequence understanding. in Advances in Neural Information Processing Systems 35 35156–35173 (Neural Information Processing Systems Foundation, Inc. (NeurIPS), San Diego, California, USA, 2022).

5. Rao, R. et al. Evaluating protein transfer learning with TAPE. Adv. Neural Inf. Process. Syst. 32, 9689–9701 (2019).

6. Walsh, I., Pollastri, G. & Tosatto, S. C. E. Correct machine learning on protein sequences: a peer-reviewing perspective. Brief. Bioinform. 17, 831–840 (2016).

7. Rives, A. et al. Biological structure and function emerge from scaling unsupervised learning to 250 million protein sequences. Proc. Natl. Acad. Sci. U. S. A. 118, e2016239118 (2021).

8. Lin, Z. et al. Evolutionary-scale prediction of atomic-level protein structure with a language model. Science 379, 1123–1130 (2023).

9. Hayes, T. et al. Simulating 500 million years of evolution with a language model. Science 387, 850– 858 (2025).

10. Stärk, H., Dallago, C., Heinzinger, M. & Rost, B. Light attention predicts protein location from the language of life. Bioinform. Adv. 1, vbab035 (2021).

11. NaderiAlizadeh, N. & Singh, R. Aggregating residue-level protein language model embeddings with optimal transport. Bioinform. Adv. 5, vbaf060 (2025).

12. Hoang, M. & Singh, M. Locality-aware pooling enhances protein language model performance across varied applications. Bioinformatics 41, i217–i226 (2025).

13. Zhang, Y. et al. DeepSecE: A deep-learning-based framework for multiclass prediction of secreted proteins in Gram-negative bacteria. Research (Wash. D.C.) 6, 0258 (2023).

14. Tang, M. et al. Predicting epistasis across proteins by structural logic. Proc. Natl. Acad. Sci. U. S. A. 123, e2516291123 (2026).

15. Jiang, X., Stockwell, B. R. & Conrad, M. Ferroptosis: mechanisms, biology and role in disease. Nat. Rev. Mol. Cell Biol. 22, 266–282 (2021).

16. Gorgoulis, V. et al. Cellular senescence: Defining a path forward. Cell 179, 813–827 (2019).

17. Muñoz-Espín, D. & Serrano, M. Cellular senescence: from physiology to pathology. Nat. Rev. Mol. Cell Biol. 15, 482–496 (2014).

18. Broz, P., Pelegrín, P. & Shao, F. The gasdermins, a protein family executing cell death and inflammation. Nat. Rev. Immunol. 20, 143–157 (2020).

19. Zhou, N. & Bao, J. FerrDb: a manually curated resource for regulators and markers of ferroptosis and ferroptosis-disease associations. Database (Oxford) 2020, baaa021 (2020).

20. Zhou, N. et al. FerrDb V2: update of the manually curated database of ferroptosis regulators and ferroptosis-disease associations. Nucleic Acids Res. 51, D571–D582 (2023).

21. Zhou, N. et al. FerrDb V3: expanding the manually curated resource for regulators and disease associations from ferroptosis to regulated cell death. Nucleic Acids Res. 54, D572–D582 (2026).

22. Lin, L. et al. FRP-XGBoost: Identification of ferroptosis-related proteins based on multi-view features. Int. J. Biol. Macromol. 262, 130180 (2024).

23. Zhou, J. & Wang, C. Enhancing ferroptosis-related protein prediction through multimodal feature integration and pre-trained language model embeddings. Algorithms 18, 465 (2025).

24. Zhao, Y. et al. Contrastive representation learning and capsule networks enable accurate identification of ferroptosis-related proteins. J. Cheminform. 18, (2026).

25. Jiang, H. et al. SenSeqNet: A deep learning framework for cellular senescence detection from protein sequences. Aging Cell 25, e70344 (2026).

26. Steinegger, M. & Söding, J. MMseqs2 enables sensitive protein sequence searching for the analysis of massive data sets. Nat. Biotechnol. 35, 1026–1028 (2017).

27. McInnes, L., Healy, J., Saul, N. & Großberger, L. UMAP: Uniform Manifold Approximation and Projection. J. Open Source Softw. 3, 861 (2018).

28. Ferrer Florensa, A., Almagro Armenteros, J. J., Nielsen, H., Aarestrup, F. M. & Clausen, P. T. L. C. SpanSeq: similarity-based sequence data splitting method for improved development and assessment of deep learning projects. NAR Genom. Bioinform. 6, lqae106 (2024).

29. Avelar, R. A. et al. A multidimensional systems biology analysis of cellular senescence in aging and disease. Genome Biol. 21, 91 (2020).

30. The UniProt Consortium et al. UniProt: the universal protein knowledgebase in 2021. Nucleic Acids Res. 49, D480–D489 (2021).

31. Vaswani, A., et al. Attention is all you need. arXiv [cs.CL] (2017) doi:10.48550/arXiv.1706.03762.

32. Loshchilov, I. & Hutter, F. Decoupled weight decay regularization. arXiv [cs.LG] (2017) doi:10.48550/arXiv.1711.05101.

33. Jain, S. & Wallace, B. C. in *Proceedings of the 2019 Conference of the North* (Association for Computational Linguistics, Stroudsburg, PA, USA, 2019). doi:10.18653/v1/n19-1357.

34. Kanehisa, M., Furumichi, M., Sato, Y., Matsuura, Y. & Ishiguro-Watanabe, M. KEGG: biological systems database as a model of the real world. Nucleic Acids Res. 53, D672–D677 (2025).

35. Milacic, M. et al. The reactome pathway knowledgebase 2024. Nucleic Acids Res. 52, D672–D678 (2024).

36. Agrawal, A. et al. WikiPathways 2024: next generation pathway database. Nucleic Acids Res. 52, D679–D689 (2024).

37. The Gene Ontology Consortium. The Gene Ontology knowledgebase in 2026. Nucleic Acids Res 54, D1779–D1792 (2026).

